# Functional and *in-silico* interrogation of rare genomic variants impacting RNA splicing for the diagnosis of genomic disorders

**DOI:** 10.1101/781088

**Authors:** Jamie M Ellingford, Huw B Thomas, Charlie Rowlands, Gavin Arno, Glenda Beaman, Beatriz Gomes-Silva, Christopher Campbell, Nicole Gossan, Claire Hardcastle, Kevin Webb, Christopher O’Callaghan, Robert A Hirst, Simon Ramsden, Elizabeth Jones, Jill Clayton-Smith, Andrew R Webster, Genomics England Research Consortium, Raymond T O’Keefe, William G Newman, Graeme CM Black

**Author notes:** These authors contributed equally (Huw B Thomas, Charlie Rowlands).

## Abstract

**Purpose:** To develop a comprehensive analysis framework to identify pre-messenger RNA splicing mutations in the context of rare disease.

**Methods:** We assessed ‘variants of uncertain significance’ through six *in-silico* prioritization strategies. Firstly, through comparison to functional analyses, we determined the precise effect on splicing of variants identified through clinical multi-disciplinary meetings. Next, we calculated the sensitivity of *in-silico* prioritization strategies to distinguish known splicing mutations from common variation (>2% in allele frequency in gnomAD) within relevant disease genes. These approaches defined an accurate *in-silico* strategy for variant prioritization, which we retrospectively applied to a large cohort of 2783 individuals who had previously received genomic testing for rare genomic disorders. We assessed the clinical impact of such prioritization strategies alongside routine diagnostic testing strategies.

**Results:** We identified 21 variants that potentially impacted splicing, and used cell based splicing assays to identify those variants which disrupted normal splicing. These findings underpinned new molecular diagnoses for 14 individuals. This process established that the use of pre-defined thresholds from a machine learning splice prediction algorithm, SpliceAI, was the most efficient method for variant prioritization, with a positive predictive value of 86%. We analysed 1,346,744 variants identified through diagnostic testing for 2783 individuals and observed that splicing variant prioritization strategies would improve clarity in clinical analysis for 15% of the individuals surveyed. Prioritized variants could provide new molecular diagnoses or provide additional support for molecular diagnosis for up to 81 individuals within our cohort.

**Conclusion:** We present an *in-silico* and functional analysis framework for the assessment of variants impacting pre-messenger RNA splicing which is applicable across monogenic disorders. Incorporation of these strategies improves clarity in diagnostic reporting, increases diagnostic yield and, with the advent of targeted treatment strategies, can directly alter patient clinical management.

**Key Highlights:**

- We establish an *in-silico* and functional analysis framework for the incorporation of splice variant assessment into diagnostic testing that is applicable across monogenic disorders.
- After assessment of six distinct variant prioritization strategies, we concluded that SpliceAI was the best method to accurately identify genomic variation disrupting normal pre-mRNA splicing. We determined this through (i) functional assessment of novel ‘variants of uncertain significance’ described in this study, and (ii) calculation of sensitivity and specificity for prioritization strategies to distinguish known splicing mutations from common variants in the general population.
- We describe novel disease-causing variants with support from cell based functional assays which underpin autosomal recessive, autosomal dominant and X-linked Mendelian disorders. This includes variants which are deeply intronic, within the nearby splice region of canonical splice sites and variants which activate cryptic splice sites within the protein-coding regions of genes.
- We integrated the best performing variant prioritization strategy alongside clinical diagnostic testing for 2783 individuals referred to a well-established targeted gene panel test available through the UK National Health Service. We show that integration of such strategies will increase accuracy and clarity of diagnostic reporting, including the identification of variants which could provide new diagnoses and new carrier findings for referred individuals.
- Functional assessment is essential for accurate clinical assessment of variants disrupting pre-mRNA splicing. We show through cell based functional assessments that variants impacting splicing may have complex impacts on pre-mRNA splicing, which may cause multiple interpretable consequences according to ACMG guidelines.

## Introduction

Pinpointing disease-causing genomic variation informs diagnosis, treatment and management for a wide range of rare disorders. Molecular testing, in a healthcare setting, now frequently includes complete genomic and exome sequencing.^1–3^ Accurate interpretation and categorization of identified variants remains a key limiting factor despite the availability of guidelines for variant analysis.^4,5^

With the advent of whole genome sequencing healthcare strategies,^3,6–9^ there is an opportunity for diagnostic services to routinely include the analysis of non-coding variation to increase diagnostic yields and improve patient care. Such analysis strategies may include consideration of variants within characterized regulatory elements,^10–12^ variants disrupting chromatin conformation^13,14^ and variants that disrupt vital processes during gene expression, e.g. pre-mRNA splicing.^15,16^ However, the capability to interpret variation within the non-coding genome is particularly challenging. Variant interpretation is hindered by the vast number of rare/novel non-coding variants identified in each individual,^7,9^ the depleted levels of evolutionary conservation within non-coding regions,^17^ and our current lack of understanding of the motifs and interactions that are required for appropriate control of gene expression and regulation.^12,18^

Intragenic genomic variants have the potential to impact splicing,^16^ the ubiquitous process in eukaryotic cells of converting nascent pre-mRNA molecules into a mature messenger RNA (mRNA) which can be transported out of the nucleus to provide a template for protein synthesis. Genomic variation in protein-coding, splice junction and intronic regions of genes can disrupt normal splicing mechanisms and underpin the onset of rare disease.^19^ Known mechanisms of splicing disruption include the introduction of cryptic splice sites, disruption of canonical splice acceptor and donor sites, and the disruption of motifs essential for splicing, e.g. branch points and the polypyrimidine tract.^19^ A number of computational tools have been developed to assist in the interpretation of genomic variation impacting splicing, and these tools have been expanded recently to include an array of machine learning tools that have been trained to prioritize splice disrupting variation through diverse means.^20–24^ While the initial reports of these tools have shown promising results there is yet to be a formal assessment of their integration, utilization and comparative performance in clinical environments.

In this study we interrogate data derived from targeted gene panel sequencing and whole genome sequencing from individuals with a broad spectrum of rare disorders. We apply *in-silico* splicing tools to prioritize variants predicted to impact splicing and we assess variant pathogenicity using targeted cDNA amplification and cell based minigene assays. We establish methodologies to identify novel causes of autosomal recessive, autosomal dominant and X-linked genomic disorders and demonstrate a requirement for routine diagnostic services to assess the effect of protein-coding and non-coding variation on mRNA expression.

## Methods

### Patient Recruitment & Genomic Variant Dataset Generation

All individuals included in this study have provided consent for the analysis of relevant disease causing genes through tertiary healthcare centers within the UK. All individuals with whole genome sequencing datasets have consented through the Genomics England 100,000 Genomes Project.

For real-time assessment of variant prioritization strategies, we identified individuals with ‘variants of uncertain significance’ according to ACMG guidelines for variant interpretation.^4^ In all cases we considered inheritance modes associated with monogenic disorders available in OMIM (https://omim.org/) or PanelApp (https://panelapp.genomicsengland.co.uk/), the zygosity of identified variants, additional variants identified to impact the same gene, phenotype-genotype correlations and scores determined by *in-silico* splicing tools (Figure 1).

**Figure 1.**
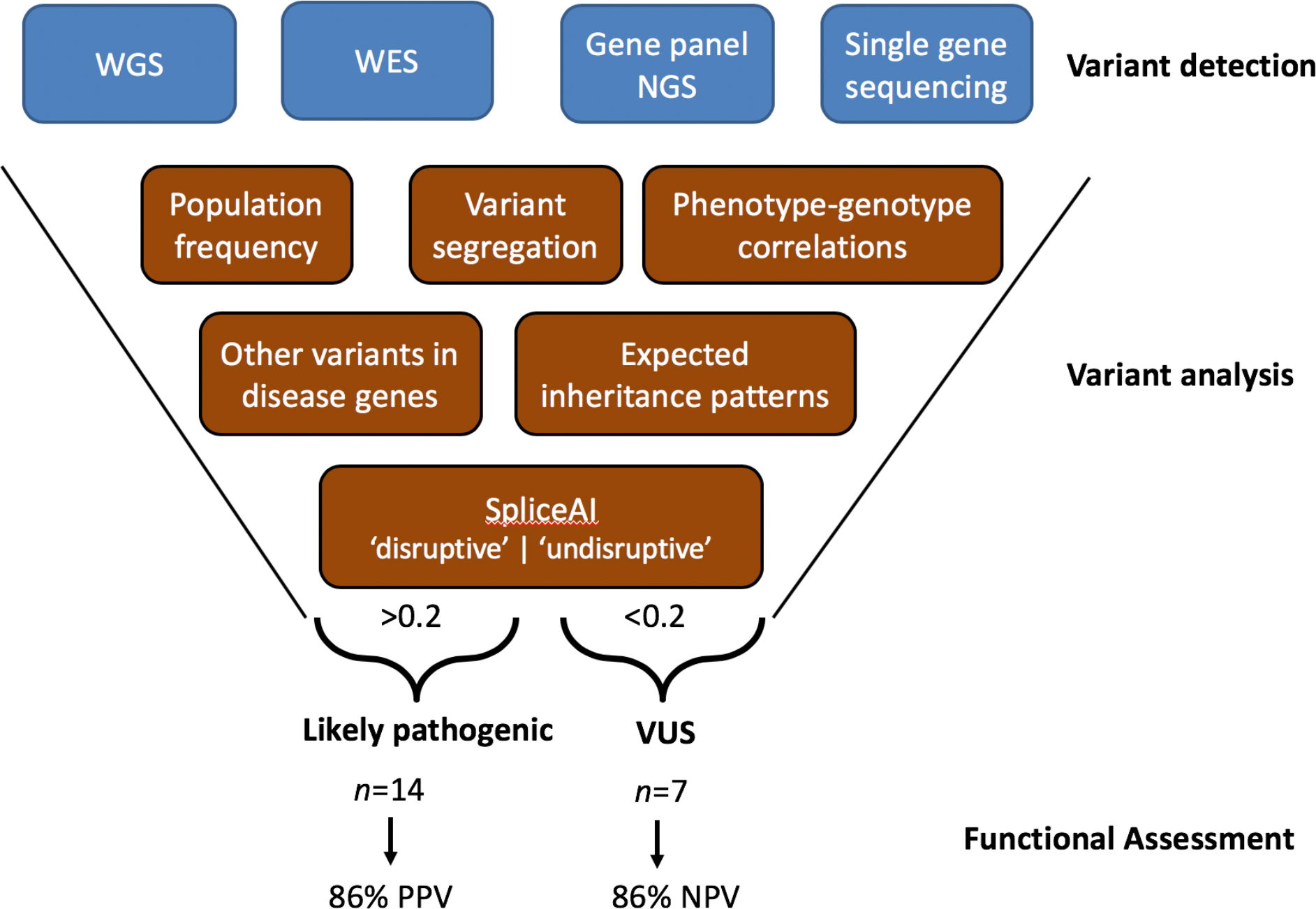
Variant prioritization strategy overview, illustrating the best performing strategy to prioritize variants expected to impact splicing. *WGS*, whole genome sequencing; *WES*, whole exome sequencing; *NGS*, next-generation sequencing; *PPV*; positive predictive value; *NPV*; negative predictive value.

#### Whole genome sequencing

Whole genome sequencing datasets were created through the UK 100,000 genomes project,^3^ using Illumina X10 sequencing chemistry. Sequencing reads were aligned to build GRCh37 of the human reference genome utilizing Issac. Small variants were identified through Starline (SNV and small indels ≤ 50bp), and structural variants were identified utilizing Manta and Canvas (CNV Caller). Variants were annotated and analysed with the Ensembl variant effect predictor (v92), bcftools and bespoke perl scripts within the Genomics England secure research embassy.

#### Gene panel sequencing

Enrichments were performed on DNA extracted from peripheral blood using Agilent SureSelect Custom Design target-enrichment kits (Agilent, Santa Clara, CA, USA). Enrichment kits were designed to capture known pathogenic intronic variants and the protein-coding regions +/−50 nucleotides of selected NCBI RefSeq transcripts; conditions tested included inherited retinal disease (105 genes or 176 genes), ophthalmic disorders (114 genes), cardiac disorders (72 genes comprised of 10 sub-panels) and severe learning difficulties (82 genes). Genes tested and relevant testing strategies are available through the UK Genetic Testing Network (https://ukgtn.nhs.uk/). All samples included in the large cohort analysis were generated through a previously described methodology,^25^ and had been completed prior to August 2017. Briefly, samples were pooled and paired-end sequencing was performed using the manufacturer protocols for the Illumina HiSeq 2000/2500 platform (Illumina, Inc., San Diego, CA, USA). Sequencing reads were demultiplexed with CASAVA v.1.8.2. and aligned to the GRCh37 reference genome using Burrows-Wheeler Aligner short read (BWA-short v0.6.2) software before duplicate reads were removed using samtools v0.1.18. The detection and clinical analysis of single nucleotide variants and small insertions/deletions was performed as described previously,^25,26^ and in accordance with ACMG guidelines for variant interpretation.^4^

### *In-silico* splicing prediction scores

#### Real-time assessment of variant prioritization strategies

*In-silico* splicing prediction scores were generated through the latest web server for the algorithm, through access to pre-computed scores, or through 3^rd^ party software (Ensembl Variant Effect Predictor or Alamut Visual Software). We utilized scores available from the following algorithms: CADD,^27^ SpliceAI,^20^ SPIDEX,^24^ S-CAP^22^ and MaxEntScan.^28^ Where multiple scores were available for a variant from the *in-silico* tool, we selected the highest score for consideration. Where scores were unavailable we arbitrarily assigned the variant a score of 0. Pre-defined thresholds were applied to determine whether a variant was ‘disruptive’ or ‘undisruptive’ to splicing, as suggested by the authors of the original papers,^20,24^ by recent refinements of thresholds,^22^ or through nationally recommended guidelines.

#### Assessment of variant prioritization strategies for known disease genes

To determine the performance of three *in-silico* splicing tools (CADD, SPIDEX and SpliceAI) within genes known as a cause of inherited retinal disease, we identified sets of variants which could be considered true negatives (no expected impact on splicing) or true positives (expected impact of splicing). Our genes of interest are listed in Table S1. True negative (TN) variants were defined as variants available through the gnomAD web server with an allele frequency above 2%. True positive (TP) variants were defined as ‘splicing’ variants available through HGMD professional. The TN and TP variant datasets were intersected and any overlapping variants were removed from subsequent analysis. For each of the *in-silico* tools, we analysed TN and TP variants using the ‘pROC’ R package to calculate the area under the curve (AUC) and to calculate tool-specific thresholds for the optimal separation of TN and TP variants.

### RNA investigations

Appropriate functional assays were selected after consideration of gene expression profiles in GTEX (https://gtexportal.org/home/), and the availability of relevant patient samples.

#### RNA investigations from patient samples – LCLs and blood

Lymphoblast cell cultures were established for control samples and probands. RNA was extracted using the RNeasy^®^ Mini Kit (Qiagen, UK, Catalogue No. 74104) following the manufacturer’s protocol. RNA was extracted from whole-cell blood using the PAXgene™ Blood RNA System Kit (Qiagen, UK. Catalogue No. 762174), following the manufacturer’s protocol for control samples and probands. Extracted RNA was reverse transcribed using the High Capacity RNA to cDNA Kit (Applied Biosystems, UK. Catalogue No. 4387406) following the manufacturer’s protocol. Gene specific primers (available on request) amplified relevant regions of the genes being investigated. PCR products were visualized on an agarose gel using a BioRad Universal Hood II and the Agilent 2200 Tapestation. Visualized bands were cut out and prepared for capillary sequencing on an ABI 3730xl DNA Analyzer.

#### RNA investigations using cell based minigene assays

Assays were designed to amplify appropriate genomic regions from patient DNA templates. For variants nearby to wild-type exons, we amplified regions containing one or multiple exons along with flanking ~200 intronic nucleotides. For deeply intronic variants we amplified regions containing at least 500bp of flanking intronic sequence. Primer sequences are available upon request. All regions were amplified from patient DNA templates. For homozygous variants, we also generated a minigene plasmid from a control DNA template. Amplified fragments were checked for size using gel electrophoresis, purified using the QIAquick Gel Extraction kit (Qiagen, UK, Catalogue No. 28706) and then cloned into a customized minigene plasmid (a derivative of the pSpliceExpress vector)^29^ containing an RSV-promoter and two control exons (rat insulin exons 2 and 3) using the NEBuilder^®^ HiFi DNA assembly (NEB, E2621). Amplified fragments were inserted between the two control exons. Plasmids were transformed into competent bacteria (XL-1 blue) and incubated overnight at 37°C on LB plates containing Carbenicillin. Individual colonies were cultured overnight before isolation of plasmid DNA using the GenElute™ miniprep kit (Sigma-Aldrich, Catalogue No. PLN350). Purified plasmids were Sanger sequenced to confirm successful cloning, and identify plasmids containing the wild type and variant sequence. Plasmids were transiently transfected into HEK-293 cells using Lipofectamine, and incubated for up to 48h in Dulbecco’s Modified Eagle Medium (DMEM) supplemented with 10% foetal bovine serum at 37°C and 5% CO_2_.

RNA was isolated using TRI Reagent^®^ and further purified using the RNeasy Mini Kit (Qiagen, UK, Catalogue No. 74106) which included a DNase digestion step. cDNA was synthesized from up to 4μg of purified RNA using SuperScript™ reverse transcriptase (ThermoFisher Scientific, Catalogue No. 18091200) and subsequently amplified by Phusion high-fidelity polymerase (ThermoFisher Scientific, Catalogue No. F553) using primers designed to amplify all minigene transcripts. PCR products were visualized by electrophoresis on a 1-2% agarose gel and purified using the QIAquick Gel Extraction kit. Purified PCR products were Sanger sequenced and aligned to the reference sequence for the minigene vector using the SnapGene software suite and assessed for differences in splicing between wild-type and variant minigene constructs.

## Results

### Real-time Assessment of Variant Prioritization Strategies

Over a 6-month period (May 2018 – November 2018), we worked through clinical multidisciplinary meetings to identify individuals with ‘variants of uncertain significance’ that could be responsible for a presenting clinical phenotype if they impacted pre-mRNA splicing (Figure 1). We ascertained 21 individuals that fitted this criterion. The broad spectrum of clinical phenotypes for these individuals are described in Table 1, and included individuals undergoing both WGS and targeted gene panel NGS testing.

**Table 1.** Clinical indications and testing strategies for individuals with ‘variants of uncertain significance’. All variants subsequently underwent *in-silico* and functional splicing analysis.

| Variant | Gene | Clinical Diagnosis | Testing strategy |
| --- | --- | --- | --- |
| NM_000350.2:c.5584+6T>C | *ABCA4* | Macular dystrophy | Gene panel |
| NM_000180.3:c.3043+5G>A | *GUCY2D* | Leber Congenital Amaurosis | Genome |
| NM_000283.3:c.2130-15G>A | *PDE6B* | Retinitis Pigmentosa | Genome |
| NM_006343.2:c.2486+6T>A | *MERTK* | Severe rod cone dystrophy | Gene panel |
| NM_001040142.1:c.2919+3A>G | *SCN2A* | Severe Epilepsy | Gene panel |
| NM_015600.4:c.867+5G>A | *ABHD12* | Usher Syndrome | Gene panel |
| NM_005208.4:c.213C>T | *CRYBA1* | Bilateral lamellar cataracts | Gene panel |
| NM_001277115.1:c.6547-963G>A | *DNAH11* | Primary Ciliary Dyskinesia | Genome |
| NM_000492.3:c.3874-4522A>G | *CFTR* | Cystic Fibrosis | Genome |
| NM_001034853.1:c.247G>T | *RPGR* | Retinitis Pigmentosa | Gene panel |
| NM_001034853.1:c.1754-3G>C | *RPGR* | Retinitis Pigmentosa | Gene panel |
| NM_000256.3:c.1224-21A>G | *MYBPC3* | Hypertrophic Cardiomyopathy | Gene panel |
| NM_002420.5:c.899+29G>A | *TRPM1* | Congenital Stationary Night Blindness | Gene panel |
| NM_002335.2:c.1413-7T>A | *LRP5* | Familial Exudative Vitreoretinopathy | Gene panel |
| NM_015629.3:c.528-38C>T | *PRPF31* | Rod-cone dystrophy | Gene panel |
| NM_024649.4:c.592-21A>T | *BBS1* | Retinitis Pigmentosa | Gene panel |
| NM_206933.2:c.14343+36C>G | *USH2A* | Retinitis Pigmentosa | Gene panel |
| NM_001142800.1:c.6571+4558A>G | *EYS* | Retinitis Pigmentosa | Genome |
| NM_001098.2:c.526-642C>T | *ACO2* | Syndromic, global developmental delay | Genome |
| NM_020366.3:c.491-386C>T | *RPGRIP1* | Cone Dysfunction Syndrome | Genome |
| NM_006204.3:c.1072-11T>C | *PDE6C* | Cone Dysfunction Syndrome | Genome |

We calculated *in-silico* splicing scores for each of the identified variants using a variety of algorithms, and arbitrarily assigned a value of 0 for all variants where *in-silico* scores were unavailable (Table 2). We predicted variants to be ‘disruptive’ or ‘undisruptive’ according to pre-defined thresholds from each of the *in-silico* splicing tools, resulting in dissimilar outcomes across tools (Table 2). None of the identified variants were consistently scored as ‘disruptive’ across all five *in-silico* splicing algorithms. Similarly, no variants were consistently assigned scores and defined as ‘undisruptive’ across all five *in-silico* splicing algorithms. Three deeply intronic variants were consistently scored as ‘undisruptive’, but these variants were outside of the regions of consideration for SPIDEX and S-CAP. We also applied an *in-silico* consensus approach, as suggested by ACMG guidelines (*PP3*),^4^ where variants were considered as ‘disruptive’ if they exceeded the pre-defined thresholds for 3/5 of the *in-silico* splicing tools. The 3/5 consensus approach prioritized 11 of the 21 variants (Table 2).

**Table 2.**
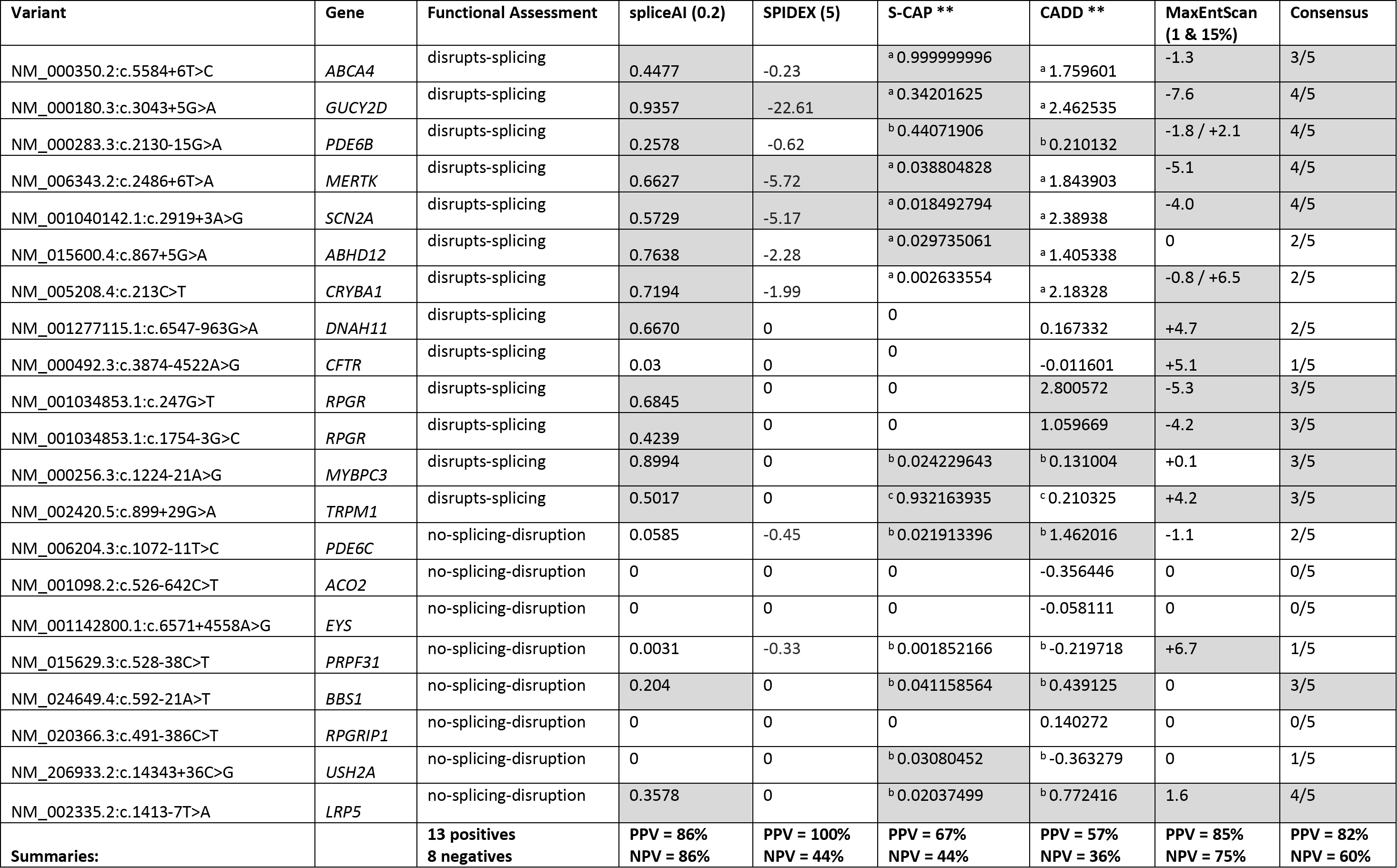
*In-silico* splicing scores calculated for ‘variants of uncertain significance’ functionally investigated to assess impact on pre-mRNA splicing. Scores unavailable from tools were arbitrary given a value of 0. The thresholds for SpliceAI was set at 0.2, and for SPIDEX at 5/−5, as per suggestions of the authors. The high-sensitivity thresholds utilized for S-CAP and CADD were categorized by region, S-CAP: ^(a)^5’extended=0.005, ^(b)^3’intronic=0.006, ^(c)^5’intronic=0.006, exonic=0.009; CADD: ^(a)^5’extended=7.39, ^(b)^3’intronic=0.0964, ^(c)^5’intronic=0.574, exonic=0.390. *Grey boxes* indicate that the variant exceeds the applied *in-silico* splicing score and was interpreted as ‘disruptive’. PPV=positive predictive value; NPV=negative predictive value.

### Functional Assessment of Prioritized Variants

To determine the accuracy of the *in-silico* splicing algorithms, we performed functional investigations to assess the exact impact of the investigated variant on splicing and predicted the effect on protein synthesis. Of the 21 variants functionally investigated, we found that 13 clearly resulted in aberrant splicing and could be reclassified as ‘likely pathogenic’ (Table 3). These findings provided new molecular diagnoses for 14 individuals.

**Table 3.**
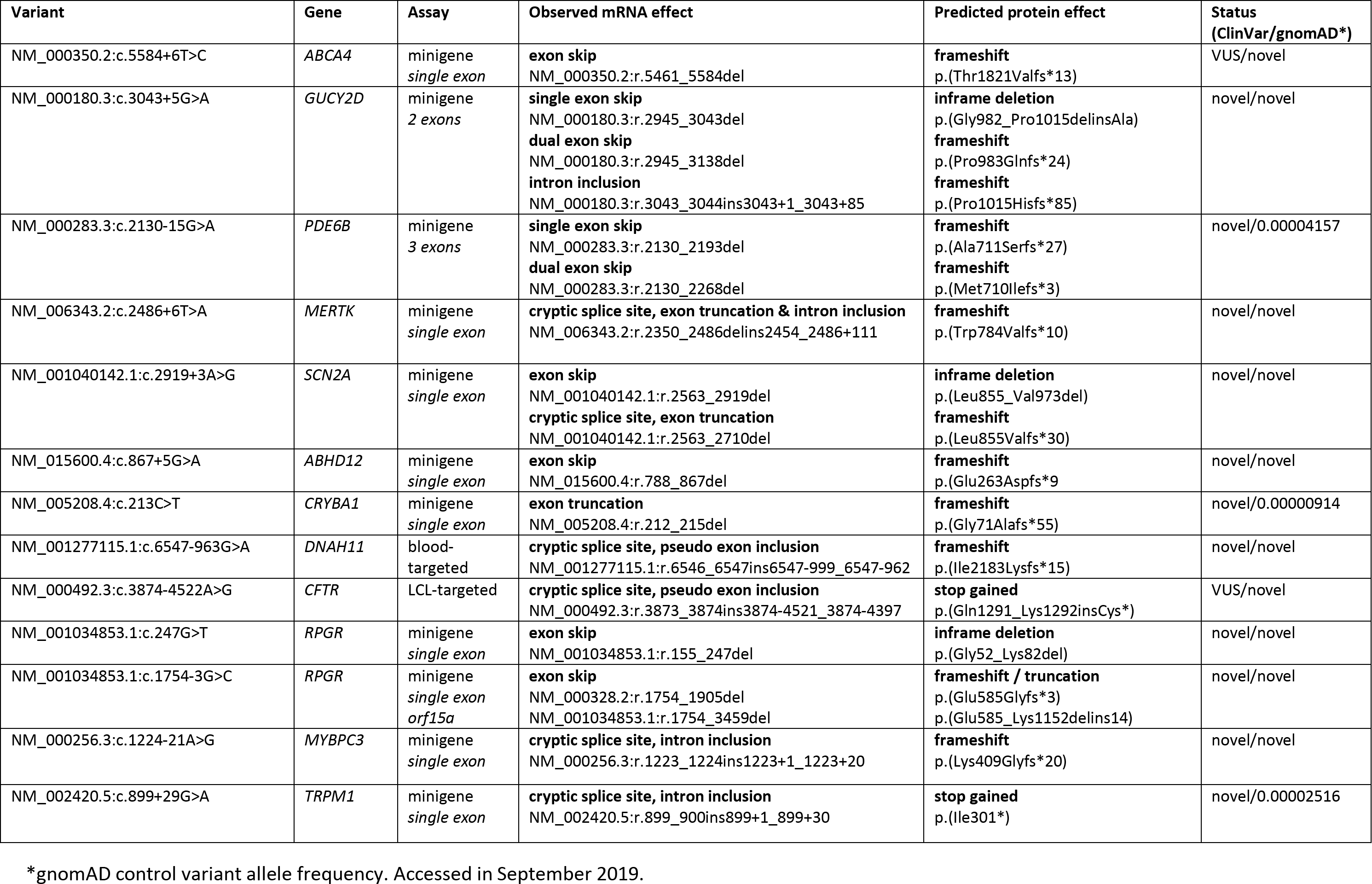
Molecular consequences of variants determined to disrupt splicing through functional assessment. The clinical phenotypes of individuals carrying these variants are described in Table 1.

| Variant | Gene | Assay | Observed mRNA effect | Predicted protein effect | Status (ClinVar/gnomAD*) |
| --- | --- | --- | --- | --- | --- |
| NM_000350.2:c.5584+6T>C | ABCA4 | minigene<br>*single exon* | **exon skip**<br>NM_000350.2:r.5461_5584del | **frameshift**<br>p.(Thr1821Valfs*13) | VUS/novel |
| NM_000180.3:c.3043+5G>A | GUCY2D | minigene<br>*2 exons* | **single exon skip**<br>NM_000180.3:r.2945_3043del<br>**dual exon skip**<br>NM_000180.3:r.2945_3138del<br>**intron inclusion**<br>NM_000180.3:r.3043_3044ins3043+1_3043+85 | **inframe deletion**<br>p.(Gly982_Pro1015delinsAla)<br>**frameshift**<br>p.(Pro983Glnfs*24)<br>**frameshift**<br>p.(Pro1015Hisfs*85) | novel/novel |
| NM_000283.3:c.2130-15G>A | PDE6B | minigene<br>*3 exons* | **single exon skip**<br>NM_000283.3:r.2130_2193del<br>**dual exon skip**<br>NM_000283.3:r.2130_2268del | **frameshift**<br>p.(Ala711Serfs*27)<br>**frameshift**<br>p.(Met710Ilefs*3) | novel/0.00004157 |
| NM_006343.2:c.2486+6T>A | MERTK | minigene<br>*single exon* | **cryptic splice site, exon truncation & intron inclusion**<br>NM_006343.2:r.2350_2486delins2454_2486+111 | **frameshift**<br>p.(Trp784Valfs*10) | novel/novel |
| NM_001040142.1:c.2919+3A>G | SCN2A | minigene<br>*single exon* | **exon skip**<br>NM_001040142.1:r.2563_2919del<br>**cryptic splice site, exon truncation**<br>NM_001040142.1:r.2563_2710del | **inframe deletion**<br>p.(Leu855_Val973del)<br>**frameshift**<br>p.(Leu855Valfs*30) | novel/novel |
| NM_015600.4:c.867+5G>A | ABHD12 | minigene<br>*single exon* | **exon skip**<br>NM_015600.4:r.788_867del | **frameshift**<br>p.(Glu263Aspfs*9) | novel/novel |
| NM_005208.4:c.213C>T | CRYBA1 | minigene<br>*single exon* | **exon truncation**<br>NM_005208.4:r.212_215del | **frameshift**<br>p.(Gly71Alafs*55) | novel/0.00000914 |
| NM_001277115.1:c.6547-963G>A | DNAH11 | blood-<br>targeted | **cryptic splice site, pseudo exon inclusion**<br>NM_001277115.1:r.6546_6547ins6547-999_6547-962 | **frameshift**<br>p.(Ile2183Lysfs*15) | novel/novel |
| NM_000492.3:c.3874-4522A>G | CFTR | LCL-targeted | **cryptic splice site, pseudo exon inclusion**<br>NM_000492.3:r.3873_3874ins3874-4521_3874-4397 | **stop gained**<br>p.(Gln1291_Lys1292insCys*) | VUS/novel |
| NM_001034853.1:c.247G>T | RPGR | minigene<br>*single exon* | **exon skip**<br>NM_001034853.1:r.155_247del | **inframe deletion**<br>p.(Gly52_Lys82del) | novel/novel |
| NM_001034853.1:c.1754-3G>C | RPGR | minigene<br>*single exon*<br>*orf15a* | **exon skip**<br>NM_000328.2:r.1754_1905del<br>NM_001034853.1:r.1754_3459del | **frameshift / truncation**<br>p.(Glu585Glyfs*3)<br>p.(Glu585_Lys1152delins14) | novel/novel |
| NM_000256.3:c.1224-21A>G | MYBPC3 | minigene<br>*single exon* | **cryptic splice site, intron inclusion**<br>NM_000256.3:r.1223_1224ins1223+1_1223+20 | **frameshift**<br>p.(Lys409Glyfs*20) | novel/novel |
| NM_002420.5:c.899+29G>A | TRPM1 | minigene<br>*single exon* | **cryptic splice site, intron inclusion**<br>NM_002420.5:r.899_900ins899+1_899+30 | **stop gained**<br>p.(Ile301*) | novel/0.00002516 |

\*gnomAD control variant allele frequency. Accessed in September 2019.

We determined positive predictive (PPV) and negative predictive values (NPV) for each *in-silico* splicing tool (Table 2). Of the assessed tools, SpliceAI outperformed the others with a PPV of 86% and a NPV of 86% (Table 2). SpliceAI prioritized 12/13 variants shown to disrupt splicing through functional assays.

We compared the use of defined thresholds from single *in-silico* splicing tools to a 3/5 consensus approach and observed that both SpliceAI and MaxEntScan outperformed the consensus approach with regards to both PPV and NPV (Table 2) for the 21 investigated variants. The consensus approach prioritized 9/13 variants shown to disrupt splicing through functional assays.

#### Cell based functional analysis enables accurate delineation of precise consequences of variants on the protein translation reading frame

We identified a variety of consequences as a result of disruption to splicing (Table 3), including variants resulting in complete exon skipping (*n*=7), and cryptic splice site activation leading to partial exon truncation (*n*=3), intron inclusion (*n*=4) and pseudo exon inclusion (*n*=2; Box 1, Case Example).

Four variants were indicated through cell based assays to have complex impacts on splicing, resulting in multiple interpretable consequences as a result of splicing disruption (Table 3). For example, *SCN2A* c.2919+3A>G produced two transcripts expressed at equal levels (Figure 2a). The first resulted in a transcript with a truncated exon, NM_001040142.1:r.2563_2710del, and the second resulted in a complete exon skip, NM_001040142.1:r.2563_2919del. While we interpreted both events as ‘likely pathogenic’ it is noteworthy that these events were considered differently using ACMG criteria;^4^ the exon truncation event resulted in a frameshift and introduction of a premature stop codon (*PVS1*), whereas the complete exon skipping event resulted in the inframe removal of 119 amino acids from the transcript (*PM4*).

**Figure 2.**
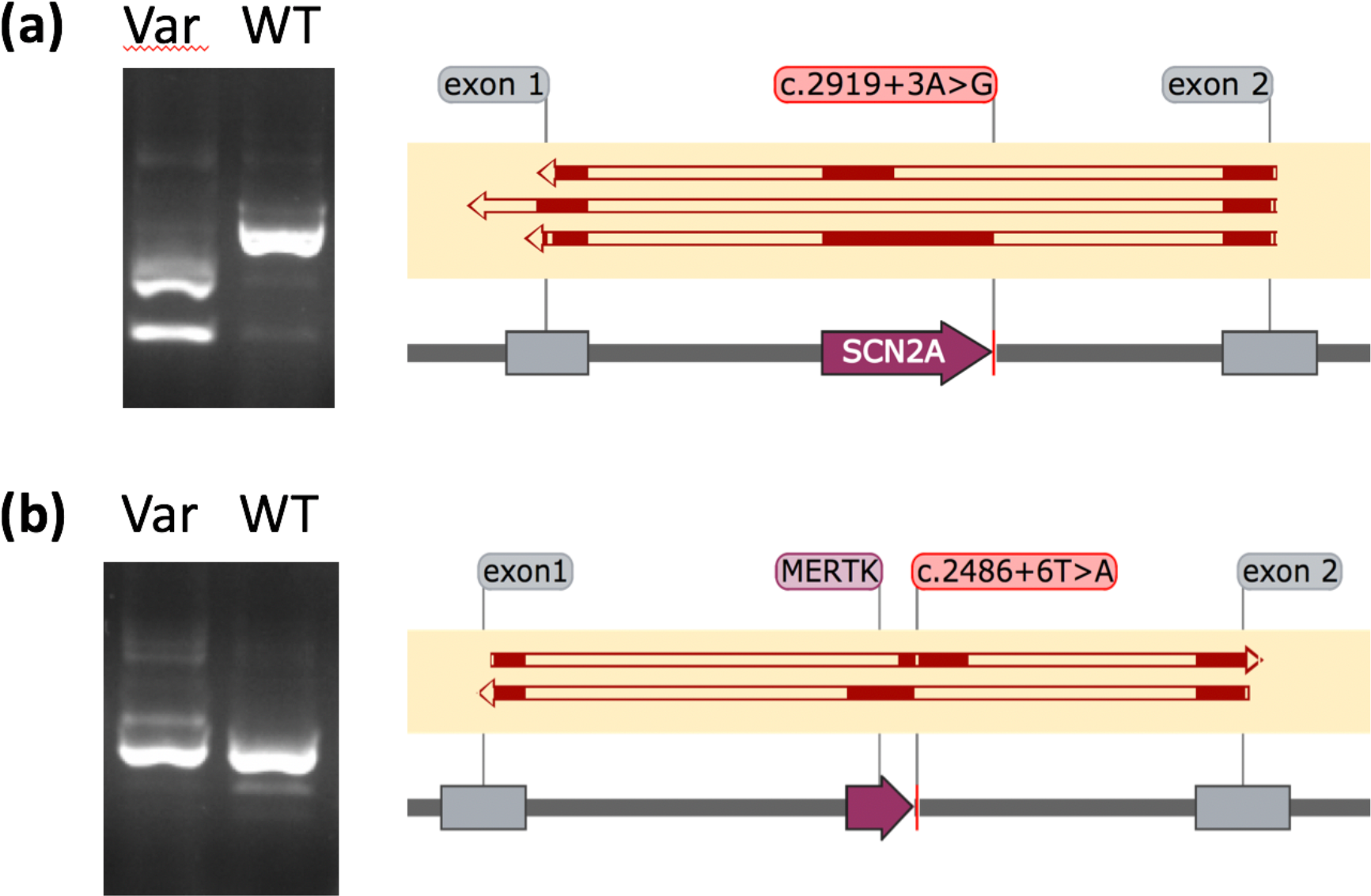
Results from *in-vitro* minigene assays demonstrating multiple consequences as a result of variants proximal to the canonical splice site. *Left*, gel electrophoresis snapshots of cDNA products amplified from primers designed for contol exons within the minigene (*exon 1* & *exon 2*). All prominent bands were cut out and Sanger sequenced. *Right*, solid red blocks illustrate alignment of sequenced cDNA transcripts to features within the minigene vector: control exons (*grey boxes*) and inserted exons (*purple boxes*). **(a) SCN2A c.2919+3A>G**, showing complete exon exclusion and exon truncation in minigene vectors containing the c.2919+3A>G variant (*top* two alignments) and normal splicing in minigene vectors containing the WT sequence (*bottom* alignment). **(b) MERTK c.2486+6T>A**, showing a shifting of the exon included in the reading frame in minigene vectors containing the c.2486+6T>A variant (*top* alignment) and normal splicing in minigene vectors containing the WT sequence (*bottom* alignment).

We identified a single instance where a variant caused usage of two new cryptic splice sites, *MERTK* c.2486+6T>A (Figure 2b). This novel variant is present in two individuals with severe rod-cone dystrophy, and resulted in the simultaneous usage of a cryptic exonic splice acceptor site and a cryptic intronic splice donor site creating a novel exon (chr2: 112,779,939-112,780,082, *GRCh37*), and a premature stop codon in the penultimate exon, p.(Trp784Valfs*10).

### Variant Prioritization Within a Large Cohort of Individuals Referred for Targeted NGS Gene Panel Testing

To evaluate the utility of *in-silico* splicing tools in a clinical environment we assessed variants identified through a targeted gene panel diagnostic test for individuals with rare genomic conditions, specifically inherited retinal disorders. All 2783 individuals had received clinical testing for known disease genes (Table S1). Overall, we demonstrated that the integration of SpliceAI with standard diagnostic testing could improve clarity and/or accuracy of diagnostic testing for 15% of the individuals analyzed, resulting in new molecular diagnoses and new carrier findings.

#### *In-silico* splicing tool assessment for relevant disease genes

We assessed the sensitivity and specificity of SpliceAI, SPIDEX and CADD to distinguish TN and TP variants within genes known as a cause of inherited retinal disease. For a communal pool of 2393 variants (TN=1068, TP=1325) which could be scored by all three prediction algorithms, we show that SpliceAI outperforms both SPIDEX and CADD, with an AUC of 0.98 and an optimal score threshold of 0.15 (Figure 3c). We also show that SpliceAI outperforms CADD and SPIDEX when considering all variants with scores available for each tool (Figure 3a) and after arbitrarily defining missing variants with a score of 0 from each tool (Figure 3b).

**Figure 3.**
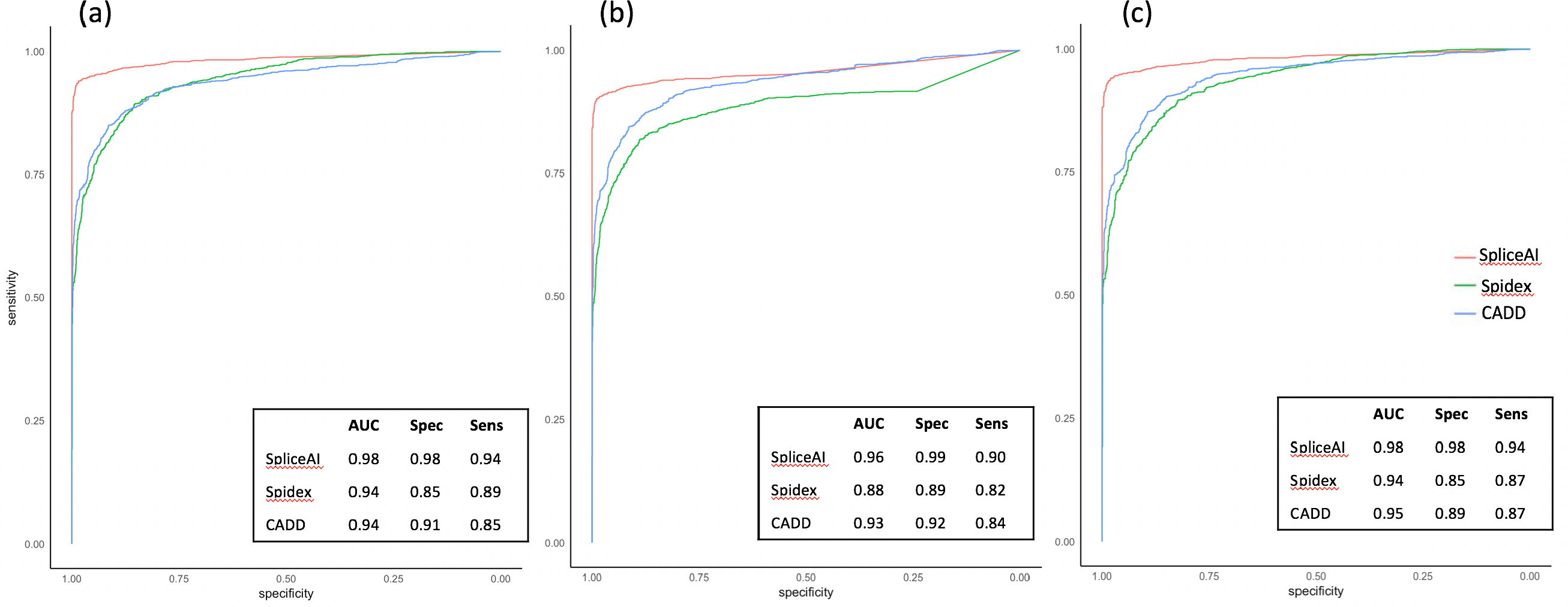
Receiver Operating Characteristic (ROC) curves comparing the performance of *in-silico* splicing tools for variants in known inherited retinal disease genes. Genes of interest are listed in TableS1. *True negatives* (TN) were defined as variants present at >2% frequency in gnomAD. *True positives* (TP) were ‘splicing’ mutations available through HGMD professional. **(a) Matched variants**, only variants which were scored by the prediction algorithm were considered (Splice AI, TN=1329, TP=1397; SPIDEX, TN=1454, TP=1335; CADD, TN=1915, TP=1436). **(b) Arbitrary values**, variants not scored by the prediction algorithm were arbitrarily scored as 0 (TN=1916, TP=1457). **(c) Communal pool**, only variants scored by all three prediction algorithms were considered (TN=1068, TP=1325). In all three situations SpliceAI outperformed SPIDEX and CADD to distinguish true negatives from true positives, with an optimal threshold score of 0.15. *AUC*, area under the curve; *Spec*, specificity; *Sens*, sensitivity.

#### Clinical impact of SpliceAI integration in variant assessment

We calculated SpliceAI scores for 1,346,744 variants (20,617 unique variants) identified in 2783 individuals. Variants exceeding a spliceAI value of 0.2 were considered alongside other variants identified as ‘pathogenic’ or ‘likely pathogenic’ after routine diagnostic testing strategies (Figure 1). We identified 758 variants (528 unique variants) in 646 individuals that were hypothesized to impact splicing of known disease genes. These variants displayed an array of predicted molecular consequences and were present in genes across disease inheritance types (Table 4; Figure 4). We defined variants prioritized through SpliceAI as:

- *New*, variant not previously highlighted or reported through diagnostic testing;
- *Clarified*, variant previously reported through diagnostic testing but pathogenicity or pathogenic mechanism was unclear;
- *Reported*, variant already described or established as ‘pathogenic’ or ‘likely pathogenic’ through diagnostic testing.

**Table 4.** Number of variants identified above specific thresholds for SpliceAI according to variant consequence and patient outcome from routine diagnostic testing strategies.

|  | Number of variants exceeding SpliceAI threshold scores: |  |  |  |  |  |
| --- | --- | --- | --- | --- | --- | --- |
| **SpliceAI score threshold:** | **>0.2** |  | **>0.5** |  | **>0.9** |  |
| Total number of variants | (758 variants) |  | (409 variants) |  | (193 variants) |  |
| **Patient status after dx testing:** | **Solved** | **Unsolved** | **Solved** | **Unsolved** | **Solved** | **Unsolved** |
| **Canonical splice site** |  |  |  |  |  |  |
| *previously reported* | 142 | 43 | 140 | 41 | 111 | 31 |
| **Splice region, synonymous, intronic** |  |  |  |  |  |  |
| *previously reported* | 30 | 11 | 18 | 9 | 1 | 3 |
| *clarified* | 34 | 16 | 27 | 11 | 6 | 2 |
| *newly identified* | 139 | 109 | 59 | 40 | 13 | 9 |
| **Missense** |  |  |  |  |  |  |
| *previously reported* | 13 | 0 | 8 | 0 | 4 | 0 |
| *clarified* | 24 | 13 | 11 | 6 | 3 | 1 |
| *newly identified* | 61 | 66 | 10 | 16 | 6 | 1 |
| **Nonsense** |  |  |  |  |  |  |
| *previously reported* | 46 | 7 | 12 | 0 | 2 | 0 |
| **UTR regions** |  |  |  |  |  |  |
| *previously reported* | 1 | 3 | 1 | 0 | 0 | 0 |

**Figure 4.**
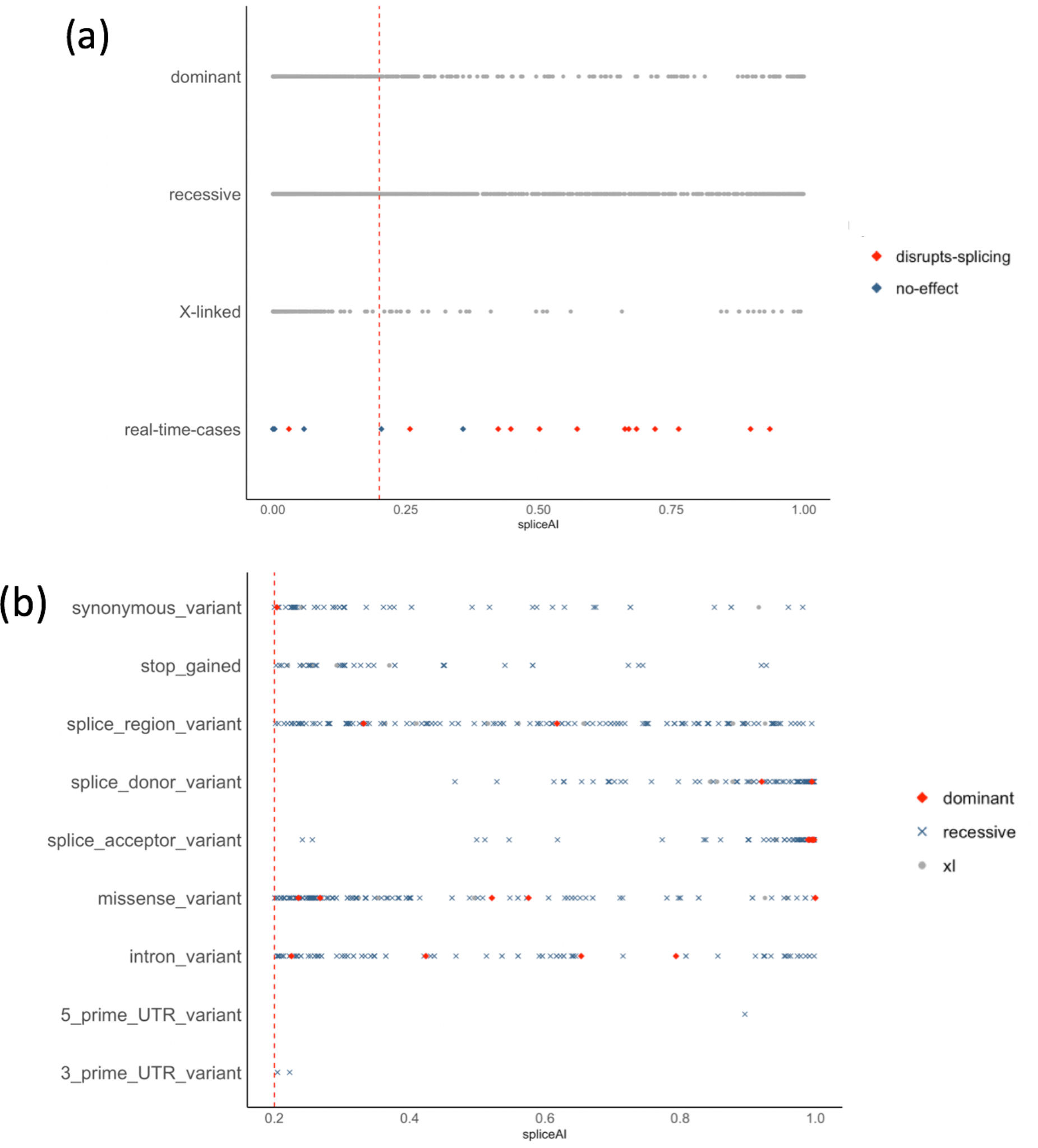
Distribution of spliceAI scores for prioritized variants identified in 2783 individuals undergoing diagnostic testing for inherited retinal disease. (a) real-time cases with functional investigations indicated, and numbers of variants within large cohort separated by inheritance patterns of associated genes. (b) distribution of scores by variant consequences. Vertical red intersect indicates the spliceAI score threshold of 0.2.

**Figure 5.**
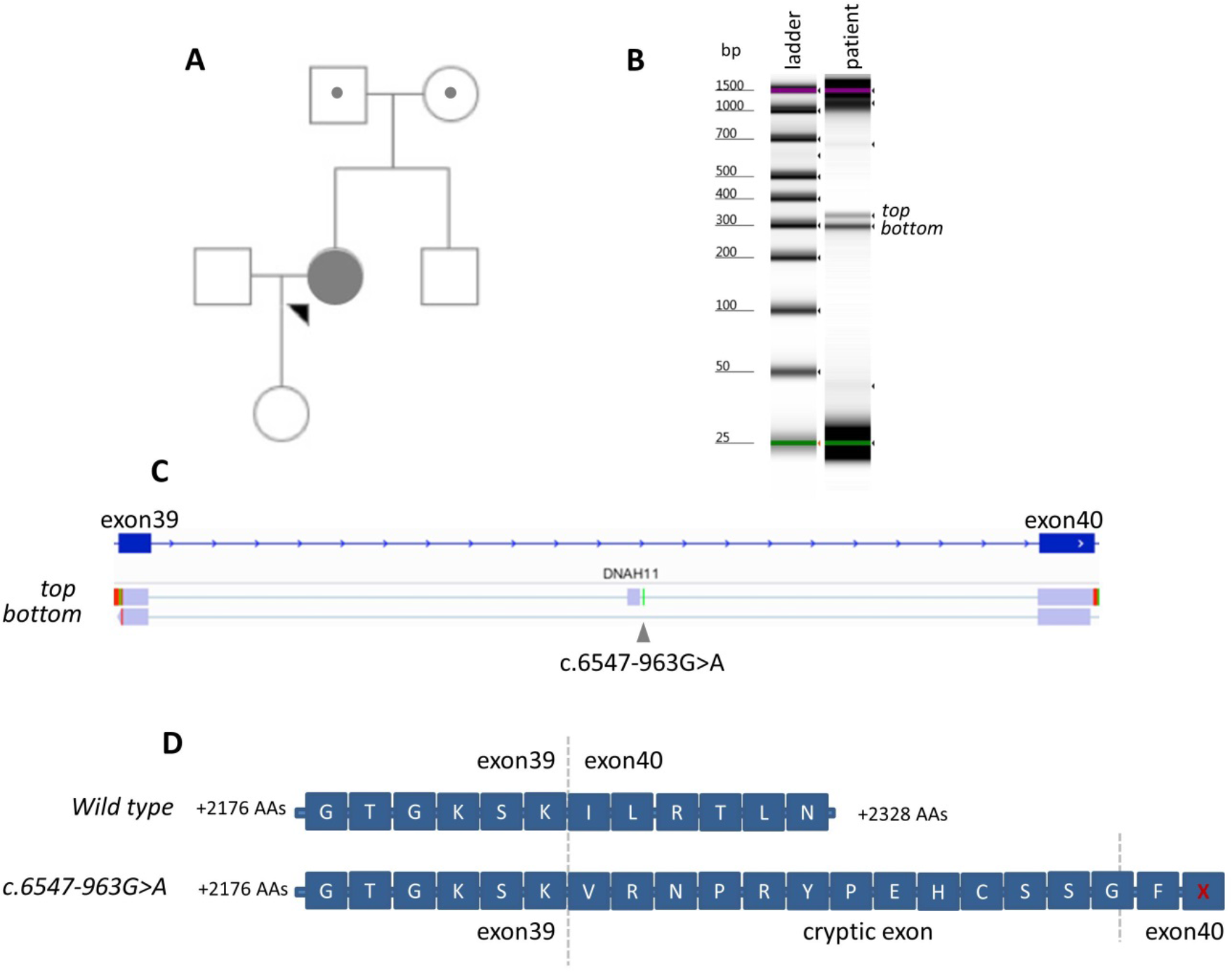
*DNAH11* c.6547-963 G>A. **(A)** Family pedigree showing the proband and her unaffected father and mother who carry heterozygous alleles of *DNAH11* c.8610C>G and c.6547-963G>A, respectively. **(B)** Gel electrophoresis results for the proband, visualized using an Agilent 2200 Tapestation. RNA was reverse transcribed after extraction from whole blood and then amplified using primers specific to exons 39 and 40 of the *DNAH11* gene (NM_001277115.1). The caption shows two distinct cDNA amplicons in the proband sample separated by ~40 base pairs. **(C)** Integrated Genomic Viewer snapshot of the alignment of sequencing products to the human reference genome (GRCh37) showing the introduction of a 38 base pair cryptic exon (chr7:21,746,318-21,746,355) as a result of c.6547-963 G>A. The *top* and *bottom* bands were sequenced after being cut from an agarose gel electrophoresis. **(D)** Impact of the cryptic exon on the translated protein. The cryptic exon shifts the reading frame and is expected to introduce a premature stop codon in exon 40, resulting in premature termination of protein synthesis, p.Ile2183Lysfs*15. Amino acids (AAs) are provided with single letter notations, with *X* indicating a stop codon. Vertical intersects indicate transition of the cDNA to the adjacent exon.

In this manner, we identified 379 *new* variants in 337 individuals, 87 *clarified* variants in 83 individuals and 292 *reported* variants in 274 individuals (Table 4). Of the 758 variants with a SpliceAI score >0.2, 145 (19%) were present in HGMD and 613 (81%) were novel. We found most (*n*=697) variants to be in genes known as a recessive cause of inherited retinal disease, variants were also prioritized in genes known as a cause of autosomal dominant (22 variants in 22 individuals) and X-linked (39 variants in 39 individuals) disorders.

#### Prioritized variants from *in-silico* splicing tools have a variety of molecular consequences

Variants with a SpliceAI score >0.2 were present in our dataset with 9 distinct predicted molecular consequences, including 185 canonical splice site variants, 100 intronic variants, 175 splice region variants, 64 synonymous variants and 177 missense variants. We segregated variants into 3 overlapping groups based on their SpliceAI score (Table 4): *above threshold*, >0.2 (*n*=758); *high*, >0.5 (*n*=409); *very high*, >0.9 (*n*=193).

In general, canonical splice site mutations had *high* SpliceAI predictions, with 181/185 variants scoring *high*, and 142/185 scoring *very high* (Table 4; Figure 4). All canonical splice site mutations prioritized through SpliceAI had been previously reported as disease-causing or carrier findings from standard diagnostic testing (Table 4).

We identified 64 synonymous, 175 splice region and 100 intronic variants above the 0.2 SpliceAI threshold. The scores for intronic, splice region and synonymous variants were wide-ranging (Table 4; Figure 4) but enabled identification of 248 potentially disease-causing or carrier mutations missed by standard testing strategies, 99 of which were identified with *high* SpliceAI scores. Of the 99 *high* scoring variants, 40 were present in individuals without a clear molecular diagnosis from diagnostic testing, and six were present in a disease-causing state. For example, we identified *high* scoring novel homozygous intronic variants in *MYO7A*, NM_000260.3:c.6559-9T>A (SpliceAI=0.9539), and in *CERKL*, NM_001030311.2:c.1238-10T>G (SpliceAI=0.8558), in individuals with Usher syndrome and retinitis pigmentosa and without clear molecular diagnoses, respectively. Ninety-three of the *high* scoring synonymous, splice region and intronic variants represented new potential carrier findings.

We identified 177 missense variants with a SpliceAI score >0.2, only 13 of which had been reported through diagnostic testing as ‘pathogenic’ or ‘likely pathogenic’ due to known disruption to splicing (Table 4). Thirty-seven of the prioritized missense variants had been clinically reported but analysis through SpliceAI highlighted or further supported that the pathogenic mechanism underpinning disease onset could be splicing disruption. 127 variants were newly prioritized missense variants, 66 of which were present in individuals without a clear molecular diagnosis after standard diagnostic testing, although only 16 of these variants had a *high* SpliceAI score. Forty-five missense variants with a score >0.2 were present in a disease-causing state, with a further 132 variants present as potential carrier findings.

#### Identification of new or clarified molecular diagnoses as a result of SpliceAI integration

We assessed the overall impact that the integration of SpliceAI would have in this large diagnostic cohort through the identification of *new* and *clarified* variants. In total we identified 466 variants in 407 individuals which had not been previously highlighted as a splicing mutation. Of the 407 individuals with new or clarified variants, 227 (56%) had clear/potential molecular diagnoses identified from diagnostic testing and 180 (44%) had not received a molecular diagnosis.

84 of the 466 *new* or *clarified* variants were present in a disease-causing state, including 9 in genes associated with X-linked disorders, 11 with dominant disorders and 64 with recessive disorders (16 homozygous, 48 compound heterozygous). These variants were present in 81 individuals, including 20 without a clear molecular diagnosis identified from diagnostic testing. In 64 individuals, the splicing variant was included on the clinical report as a potential but unconfirmed cause of disease, 29 of these variants were missense variants. 373 of the 466 variants were potential carrier findings in recessive genes, and 9 were potential carrier findings in X-linked genes.

Taken together, our analyses prioritized 758 variants in 646 individuals (23% of the cohort). 62% of the prioritized variants (466/758) were *new* or *clarified* variants, representing altered analysis of variants identified in 15% of the 2783 individuals. Such variants require functional investigation to establish precise effects on splicing and protein synthesis, but could account for new or refined molecular diagnoses in 81 individuals.

## Discussion

Interpreting splicing variants has been an active area of genetics research for decades.^16,30^ As a result, a number of motifs essential for splicing are known,^31,32^ variants disrupting the normal splicing process are well documented,^19,33^ and targeted treatments are emerging.^34,35^ However, expanding splicing mutation analysis approaches to whole genome sequencing and NGS datasets in the context of routine clinical investigation remains significantly limited. The objective of this study was to develop a clinically applicable framework for the prioritization and functional assessment of variants impacting splicing in the context of monogenic disease. In this regard, we compared the performance of a number of *in-silico* splicing tools to establish methods to efficiently prioritize variants disrupting splicing, and we validated the use of a cell based methodology for variant functional interrogation within patient clinical pathways. To assess the potential impact of incorporating variant prioritization strategies into clinically accredited diagnostic genomic testing, we expanded our *in-silico* investigations to a large cohort of individuals with rare disease. Overall, we demonstrate that utilizing defined score thresholds for a machine learning splicing prediction algorithm, SpliceAI,^20^ enables the efficient identification of variants which disrupt splicing (86% PPV), and would improve clarity in the routine clinical analysis for 15% of individuals undergoing diagnostic testing. While our analyses underpin improvements to diagnostic services and the clinical management of individuals, we encountered a number of notable limitations.

### Complexity defining appropriate test variants for comparative assessments of *in-silico* splicing tools

There are significant limitations associated with comparing *in-silico* variant prioritization approaches, including *information leakage*, a concept in machine learning where test datasets become contaminated with data from original training datasets. Information leakage inhibits the use of a pure and unbiased dataset to compare performance across tools. To overcome this obstacle we ascertained clinically relevant variants in real-time, most of which were absent from both mutation and population datasets (Table 1). While this dataset is relatively small, it enabled a fair and unbiased comparison across all of the *in-silico* tools assessed in this study. For three of the *in-silico* tools we also compared their performance for relevant disease genes, by assessing their capability to distinguish known splicing mutations from common variants. Significantly, we demonstrated that SpliceAI outperformed other *in-silico* splicing tools through both of these comparative analysis strategies, and that SpliceAI outperformed a consensus approach to variant prioritization (Table 2; Figure 3).

Another limitation of comparative approaches for splicing variants is the capability to segregate performance assessments for different categories of splicing mutations. In theory, variants could be sub-divided by their pathogenic mechanism, their effect on pre-mRNA splicing, by their predicted molecular consequence or by the location of the variant with respect to known splicing motifs. However, splicing mutations are often considered as a single class of variants and are therefore highly susceptible to the over-prioritization of *in-silico* tools to identify canonical splice site mutations, which represent the majority (~70%) of known splicing mutations.^19,24,33^ Recent studies have begun to categorize variants by location or expected effect and have demonstrated variable performances of *in-silico* splicing tools outside of the immediate splice region.^20,22^ In this study we show that while SpliceAI performs well for canonical splice site mutations (Table 4; Figure 4), we can also utilize SpliceAI as an analysis strategy to effectively prioritize intronic, missense and synonymous variants, including variants which led to new molecular diagnoses (Table 3). Interestingly, we prioritized 177 missense mutations within our large diagnostic cohort as potentially ‘disruptive’ to splicing (Table 4), reemphasizing earlier observations that splicing disruption is a significant pathogenic mechanism associated with missense mutations.^24,36–38^ Although we have been able to identify deeply intronic cryptic splice site mutations through the described techniques (Box 1, Case Example), an extensive assessment of whole genome sequencing datasets is required to comprehensively assess the success of SpliceAI and other *in-silico* splicing tools in this regard.

#### Box 1

##### Case Example

The proband (Figure 5) was diagnosed in childhood with bronchiectasis, she is currently 54 years of age. Nasal nitric oxide levels were extremely low at 4 parts per billion (ppb, normal range =<25ppb) consistent with a diagnosis of Primary Ciliary Dyskinesia (PCD).^48^ Three examinations of the proband’s cilia with electron microscopy (EM) showed a significant proportion of static and dyskinetic cilia with a high ciliary beat frequency: 20.2Hz (95%CI=19.8-20.5Hz); 22.0Hz (95%CI=20.1-23.2Hz); and 20.1Hz (95%CI=19.8-20.5Hz). EM histology showed normal dynein arms and microtubules with no ciliary disorientation, and conical ciliated protrusions were observed from epithelial cells. These findings are consistent with mutations in *DNAH11*, a recessive cause of PCD.^49^ Previous genetic testing identified a heterozygous nonsense mutation in *DNAH11*, p.Tyr2870Ter.

We generated whole genome sequencing datasets through the 100,000 genomes project. Analysis of variants was restricted to the genomic region of *DNAH11* (chr7:21,582,833-21,941,457, GRCh37) and prioritized a candidate non-coding mutation, *DNAH11* c.6547-963G>A. The variant is absent from gnomAD, ClinVar, HGMD and other samples in the 100,000 genomes project; it was determined to be a ‘variant of uncertain significance’.

We assessed *in-silico* splicing scores for *DNAH11* c.6547-963G>A, and determined that it was ‘disruptive’ according to thresholds applied to SpliceAI and MaxEntScan scores (Table 2). The variant had a *high* score from SpliceAI (0.67), but it was absent from regions of consideration for SPIDEX and S-CAP.

We performed targeted cDNA amplification from RNA extracted from whole blood (Figure 5). These analyses demonstrated aberrant splicing as a result of c.6547-963G>A, resulting in cryptic exon inclusion and the introduction of a premature stop codon, p.Ile2183Lysfs*15. The variant was determined to be ‘likely pathogenic’. Segregation analysis demonstrated that p.Tyr2870Ter and p.Ile2183Lysfs*15 are present in a compound heterozygous state in the proband (Figure 5), and were reported as the cause of PCD.

### Lessons learnt from establishing a cell based framework for the functional investigation of prioritized variants

We performed functional investigations to determine the precise impact of variants on splicing and predicted the impact on protein synthesis. In two cases we were able to directly extract RNA from patient samples. However, in the remaining 19 cases we were either unable to assess relevant RNA profiles from peripheral blood or LCLs, or unable to obtain additional samples for the referred patient. We therefore assessed the effects of prioritized variants within these regions through the utilization of minigene assays, a cell based technique which can be designed to insert bespoke genomic regions from patient DNA templates into a mammalian expression vector to assess differences in RNA transcripts produced from fragments containing wild-type and variant sequences, respectively. A limitation of minigene assays is the capability of such techniques to replicate *in-vivo* gene expression profiles. Indeed, splicing machinery may be influenced by cell-/tissue-specific factors or by genomic regions outside of the region inserted into the minigene vector, and variants may have pathogenic impacts on gene expression and/or regulation without any detrimental impact on splicing.^10,17,39^ Such factors are outside of the scope of the assays performed in this study, and therefore, whilst we calculate NPVs for each of the variant prioritization strategies, future investigations may uncover pathogenic roles for variants reported here. Despite these limitations, we have validated the use of a customizable minigene assay within real-time clinical investigations and successfully identified variants causing aberrant splicing. In the absence of appropriate patient samples for RNA investigation, this is a critical tool for clinical variant investigation.

Significantly, we demonstrate a range of functional consequences on mRNA splicing as a result of disruption to canonical splice sites, including (i) exon skipping, (ii) intron inclusion, (iii) exon truncation (Figure 2a), (iv) displacement of the reading frame (Figure 2b), and (v) a combination of events (Table 3). These observations demonstrate that the precise effect of splicing variants is an important piece of evidence for consideration during clinical variant interpretation, and may in the future lead to refinements in the exact targeted treatment appropriate for these individuals.^34,40^ Hence, while we clearly demonstrate that *in-silico* splicing analyses can prioritize and guide the identification of new clinically-relevant variants (Table 4), functional assessment of variant effect remains an important requirement to ensure clarity in clinical reporting and appropriate patient management.

### Significant impact of incorporating SpliceAI scores into routine clinical analysis

We were able to demonstrate that 13 variants caused aberrant splicing, and in so doing, provided new molecular diagnoses for 14 individuals (Table 3). Delineating the cause of disease through novel variant interpretation strategies can lead to altered clinical management.^41^ In this study, we elucidated that *SCN2A* c.2919+3A>G underpinned the molecular diagnosis for an individual with severe epilepsy and learning difficulties (Figure 2), and as a result, modified the use of medication to control seizures in the referred individual.^42^

To establish the impact of our variant prioritization strategies in everyday clinical practice, we analyzed variants identified in a large cohort of 2783 individuals with rare ophthalmic disease displaying significant genetic and clinical heterogeneity. Overall, we prioritized 379 variants which had evaded prioritization through standard diagnostic testing strategies and provided additional support for pathogenicity for 89 variants. These findings reflected altered clinical analysis in 15% of individuals, and could lead to new or refined molecular diagnosis in up to 81 individuals. Whilst this represents a significant improvement to molecular diagnostic services, we expect that the true impact of such analysis strategies will be more pronounced. The targeted NGS approaches employed within this large cohort ignore deeply intronic regions of genes, and as shown here (Box 1, Case Example) and in other studies,^43–47^ variation within these regions can cause aberrant splicing through the production of novel cryptic exons. We expect, therefore, that extending variant prioritization approaches to large cohorts of individuals with whole genome sequencing datasets will enable the identification of clinically relevant deeply intronic variants.

In summary, the recent availability of thousands of genomic datasets within healthcare amplifies the current limitations in interpreting variation within the non-coding genome. Our findings demonstrate the opportunity to expand bioinformatics analysis to the pre-mRNA regions of known disease genes and provide immediate increases to diagnostic yield. Moreover, we demonstrate a requirement to functionally assess variant impact on pre-mRNA splicing as the delineation of the precise effects may be important considerations for variant pathogenicity. Importantly, variants which impact mRNA expression are amenable to targeted therapy, e.g. antisense oligonucleotides,^34^ and have been proven to be effective in cell lines for some of the disorders investigated here.^40,47^ The prioritization and identification of pathogenic variants impacting splicing is therefore an important consideration for diagnostic services and for the development of new targeted treatments. Overall, we present an *in-silico* and functional analysis framework for the incorporation of splice variant assessment that is applicable across monogenic disorders.

## Acknowledgments

J.M.E. is funded by a postdoctoral research fellowship from the Health Education England Genomics Education Programme (HEE GEP). The views expressed in this publication are those of the authors and not necessarily those of the HEE GEP. GA is supported by a Fight for Sight (UK) Early Career Investigator award, Moorfields Eye Charity and the National Institute for Health Research (NIHR) Biomedical Research Centre at Great Ormond Street Hospital NHS Trust.

We thank Jeremy McRae for assistance in the interpretation and access of scores for SpliceAI. This research was supported by the Manchester Biomedical Research Centre, the NIHR-Biomedical Research Centre at Moorfields Eye Hospital and the UCL Institute of Ophthalmology, RP Fighting Blindness and Fight For Sight.

This research was made possible through access to the data and findings generated by the 100,000 Genomes Project. The 100,000 Genomes Project is managed by Genomics England Limited (a wholly owned company of the Department of Health). The 100,000 Genomes Project is funded by the National Institute for Health Research and NHS England. The Wellcome Trust, Cancer Research UK and the Medical Research Council have also funded research infrastructure. The 100,000 Genomes Project uses data provided by patients and collected by the National Health Service as part of their care and support. The Genomics England Research Consortium is listed alphabetically

---

**Table S1.** HGNC approved gene symbols for 180 genes known as a cause of inherited retinal disease.

|  |  |  |  |  |  |  |  |  |
| --- | --- | --- | --- | --- | --- | --- | --- | --- |
| *ABCA4* | *C1QTNF5* | *CNGA1* | *GNAT1* | *IQCB1* | *NEK2* | *PEX7* | *RGS9* | *TEAD1* |
| *ABHD12* | *C21orf2* | *CNGA3* | *GNAT2* | *ITM2B* | *NMNAT1* | *PHYH* | *RHO* | *TIMP3* |
| *ACBD5* | *C2orf71* | *CNGB1* | *GNPTG* | *KCNJ13* | *NPHP1* | *PITPNM3* | *RIMS1* | *TMEM237* |
| *ADAM9* | *C8orf37* | *CNGB3* | *GPR125* | *KCNV2* | *NPHP3* | *PLA2G5* | *RLBP1* | *TOPORS* |
| *ADAMTS18* | *CA4* | *CNNM4* | *GPR179* | *KIAA1549* | *NPHP4* | *PRCD* | *ROM1* | *TRIM32* |
| *AHI1* | *CABP4* | *CRB1* | *GPR98* | *KIF11* | *NR2E3* | *PROM1* | *RP1* | *TRPM1* |
| *AIPL1* | *CACNA1F* | *CRX* | *GRK1* | *KLHL7* | *NRL* | *PRPF3* | *RP1L1* | *TSPAN12* |
| *ALMS1* | *CACNA2D4* | *CSPP1* | *GRM6* | *LCA5* | *NYX* | *PRPF31* | *RP2* | *TTC8* |
| *ARL2BP* | *CAPN5* | *CYP4V2* | *GUCA1A* | *LRAT* | *OAT* | *PRPF4* | *RP9* | *TUB* |
| *ARL6* | *CC2D2A* | *DFNB31* | *GUCA1B* | *LRIT3* | *OFD1* | *PRPF6* | *RPE65* | *TULP1* |
| *BBIP1* | *CDH23* | *DHDDS* | *GUCY2D* | *LRP5* | *OTX2* | *PRPF8* | *RPGR* | *UNC119* |
| *BBS1* | *CDH3* | *DTHD1* | *HARS* | *LZTFL1* | *PANK2* | *PRPH2* | *RPGRIP1* | *USH1C* |
| *BBS10* | *CDHR1* | *EFEMP1* | *HMX1* | *MAK* | *PCDH15* | *RAB28* | *RPGRIP1L* | *USH1G* |
| *BBS12* | *CEP164* | *ELOVL4* | *IDH3B* | *MERTK* | *PCYT1A* | *RAX2* | *RS1* | *USH2A* |
| *BBS2* | *CEP290* | *EMC1* | *IFT140* | *MFRP* | *PDE6A* | *RBP3* | *SAG* | *VCAN* |
| *BBS4* | *CERKL* | *EYS* | *IMPDH1* | *MKKS* | *PDE6B* | *RBP4* | *SDCCAG8* | *VPS13B* |
| *BBS5* | *CHM* | *FAM161A* | *IMPG1* | *MKS1* | *PDE6C* | *RD3* | *SEMA4A* | *WDPCP* |
| *BBS7* | *CIB2* | *FLVCR1* | *IMPG2* | *MVK* | *PDE6G* | *RDH12* | *SLC24A1* | *WDR19* |
| *BBS9* | *CLN3* | *FSCN2* | *INPP5E* | *MYO7A* | *PEX1* | *RDH5* | *SNRNP200* | *ZNF423* |
| *BEST1* | *CLRN1* | *FZD4* | *INVS* | *NDP* | *PEX2* | *RGR* | *SPATA7* | *ZNF513* |

